# Odor source localization behavior of an insect enhanced by intermittent intake strategy

**DOI:** 10.1101/2024.08.26.609633

**Authors:** Shunsuke Shigaki, Takumi Matsushita, Hirono Ohashi, Noriyasu Ando, Koh Hosoda

## Abstract

This study investigated odor acquisition strategies to enhance odor plume tracking performance. Efficient odor plume tracking is a crucial ability for organisms, affecting their survival, including for insects with relatively simple nervous systems. Insects can use odor cues to locate food sources or potential mates. Odors released from a source disperse in complex patterns owing to air currents and collisions with objects, making their spread unpredictable. Organisms must therefore engage in active odor acquisition behaviors to effectively gather spatial information from this highly uncertain odor environment. This study focused on odor acquisition via wing flapping in a male silk moth and its relationship with female localization. Given the difficulty of directly intervening in wing flapping, we employed an insect-mounted robotic system to engineer interventions and investigate the relationship between wing-flapping-induced odor acquisition and localization. We found that the difference between air inflow and stoppage in odor attraction was large, and that the odor plume tracking performance was highest at 10 Hz, where odor attraction can be performed at high frequencies. Although constant strong odor acquisition improves localization performance, it increases the likelihood of movement in directions other than that of the odor source. This suggests that periodic wing flapping helps to suppress undesired movements.

## 1. Introduction

Odors with superior diffusion and persistence serve as crucial signals for communication and navigation in living organisms [1]. Consequently, the ability to efficiently track odors is an important skill directly tied to survival. This applies equally to insects, which possess relatively simple nervous systems. Insects use odors to locate feeding sites and potential mating partners [2]. However, target molecules released from odor sources mix with other molecules and are transported by turbulent airflows, creating complex spatial structures [3]. Additionally, environmental factors such as air currents and obstacles significantly influence odor diffusion, making it highly challenging to appropriately gather information from the environment and navigate toward the odor source. Nevertheless, organisms can track odors adeptly, even in complex environments, by appropriately adjusting their behavior based on the information obtained from multiple sensory organs and the quantity of odor information acquired [4,5].

The problem of tracking an odor and identifying its source is generally referred to as the odor source localization (OSL) problem [6]. OSL has the potential to address various societal needs such as identifying gas or hazardous material leaks in situations where visual exploration is difficult, detecting explosives, and locating survivors in disaster scenarios. Consequently, research has been conducted on implementing OSL capabilities in autonomous robots [7–9]. Given that OSL is a navigational problem, considerable emphasis has traditionally been placed on motion planning or search algorithms following sensory data acquisition. However, as mentioned previously, because odors are unpredictable signals, there is no guarantee that the required information can always be obtained. Some research has aimed to mimic biological odor-tracking behaviors to endow robots with odor source localization capabilities; even those with a certain degree of search performance tend to fall short compared to biological capabilities [10,11]. Therefore, acquiring accurate odor information from the environment is crucial for effective odor tracking.

Observations of various organisms have revealed that they exhibit behaviors during odor tracking that are actively aimed at collecting odors. For instance, dogs transition to a sniffing mode, which is characterized by a specific breathing pattern for odor detection [12] and crayfish manipulate water flow by ejecting jets to change the surrounding environment [13]. Similarly, the silk moth, an insect, performs odor plume tracking through vigorous wing flapping, despite its inability to fly [14]. Although the behaviors exhibited by different species vary, it has been suggested that these organisms modify the surrounding environment to facilitate effective odor plume tracking. This implies that high performance in odor tracking is likely a result of not only exceptional search behavior strategies, but also a combination of odor acquisition strategies. By elucidating the relationship between search behavior and odor acquisition strategies, we can gain insight into how organisms handle odors with high uncertainty and low spatiotemporal resolution. Furthermore, replicating this relationship in an engineering context is expected to lead to the development of autonomous robots with enhanced odor-plume tracking capabilities.

Therefore, this study aimed to investigate an active odor acquisition strategy that enhances the odor plume tracking performance of organisms. In this study, using a male silk moth, known for its specific response to sex pheromones released by females and its resultant odor plume tracking behavior[15], we investigated the relationship between odor acquisition strategies and odor tracking behavior. The silk moth is known to actively acquire odors by vigorously flapping its wings during female localization, and the localization performance of the silk moth decreases when the wings are removed [14]. We employed an artificial system capable of intervening in the odor acquisition strategy to investigate the relationship between various conditions of odor acquisition strategies and odor plume tracking performance, because wing removal alone has limitations in modifying active odor acquisition strategies.

## 2. Problem statement

This study investigated the relationship between odor acquisition and odor plume tracking behavior in silk moths using an artificial system capable of intervening in odor acquisition strategies. The silk moth responds to the sex pheromone (bombykol) of conspecific females, which triggers female localization behavior. During this process, the male silk moth actively flutters its wings and walks towards the female. This wing-flapping behavior not only provides propulsion[16] but also propels odors towards itself[17]. Moreover, the frequency of wing flapping is not constant during female localization, varying between 20 and 40 Hz[17,18]. In this study, we refer to the effect of the moth propelling odors towards itself through wing-flapping as the “flapping effect.”

It is hypothesized that organisms including, but not limited to, the silk moth, demonstrate superior exploratory abilities by performing intermittent inhalation, which reduces sensory adaptation to odors. By varying the frequency of inhalation, it is expected that the intervals of odor input to the sensory organs change, and performing inhalation at an optimal frequency enhances odor plume tracking performance. Accordingly, this study focuses on the intermittent odor acquisition induced by the flapping effect and elucidates its relationship with odor plume tracking behavior.

To investigate the relationship between intermittent odor acquisition and odor plume tracking behavior, we need to intervene in the flapping effect. However, accessing the flight muscles of the silk moth and arbitrarily setting the wing-flapping frequency presents significant challenges. Hence, we proposed addressing this issue by designing a device that generates a simulated flapping effect and intermittently presents odors to the silk moth at selectable frequencies. Conducting experiments with free-walking moths was not feasible using this device. Therefore, we utilized an insect-mounted robotic system[19,20] to explore the relationship between intermittent odor acquisition and tracking behavior. Specifically, we designed a robotic system to satisfy the following requirements:

1. the system can intermittently obtain odors at any frequency and provide them to the silk moth,
2. the system implements measurement and drive systems that are capable of detecting changes in the silk moth’s behavior in real time and realizing odor plume tracking.

The device used for intermittent odor acquisition is referred to as an artificial odor acquisition (AOA) device. The robotic system was engineered to detect the movement of a silk moth and drive the robot with a delay time of less than 0.2 seconds. This design consideration is based on reports indicating that the localization performance of the silk moth deteriorates when the delay exceeds 0.2 seconds in similar insect-mounted robotic systems. Odorplume tracking experiments were conducted using this insect-mounted robotic system, and behavioral changes were comprehensively measured across various acquisition frequencies. In these experiments, we physically removed the wings of the silk moth to eliminate the flapping effect.

## 3. Construction of an intervention system

### 3.1. System overview

The outline and configuration of the insect-mounted robotic system are illustrated in Figure 1. The system is equipped with an artificial odor acquisition (AOA) device that mimics the function of wing flapping and is mounted in front of the silk moth. The silk moth has two antennae on its head, so the AOA device is positioned in front of these antennae. The behavioral measurement system within the insect-mounted robotic system employs a tether. As shown in Figure 1, we secured the back of the silk moth to the fixture and positioned it on a passively rotating sphere. The rotation of the sphere is detected by optical sensors that convert the rotational data into movement data to control the drive system. These operations are performed using a microcontroller (Arduino Mega). A Bluetooth module attached to the microcontroller wirelessly transmits data, such as the movement data of the silk moth, to a computer for logging.

**Figure 1.**
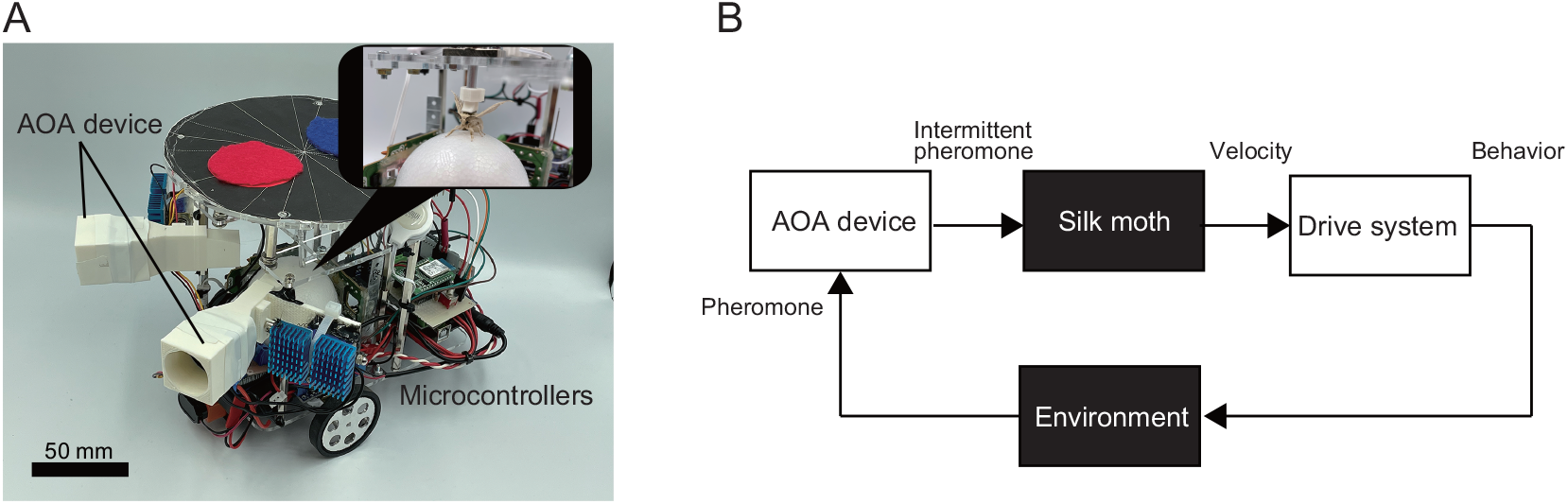
Outline of the insect-mounted robot system and block diagram of the system.

### 3.2. Artificial odor acquisition device

An overview of an artificial odor acquisition (AOA) device that intermittently measures odors at arbitrary frequencies is illustrated in Figure 2A. The AOA device incorporated a small fan (3010H12S, Wide Work Corporation, Tokyo, Japan) installed at the air intake and a gate valve positioned immediately before the air outlet to achieve intermittent airflow. The gate valve was controlled using a solenoid (JF-0730B-12V, Fielect, China) to enable initiation and cessation of airflow at the desired intervals. The solenoid was equipped with a spring; when voltage was applied, the magnetic force of the solenoid lifted the gate valve, and when the voltage was released, the spring force caused the gate valve to descend. Voltage control was achieved using an N-channel MOSFET (2SK2232; TOSHIBA Corporation, Tokyo, Japan). A pulse voltage with a 50% duty cycle was applied at various frequencies to regulate the opening and closing of the gate valve. For instance, controlling the gate valve at a frequency of 1 Hz involved setting open and closed durations of 0.5 seconds each. The detailed airflow path was designed as shown in the three-view diagram (Figure 2B), and manufactured using a 3D printer (Creator3, Flashforge Technology Co., Ltd., Zhejiang, China). Additionally, a heat sink was installed to dissipate heat because the solenoid became very hot when operated continuously for long periods.

**Figure 2.**
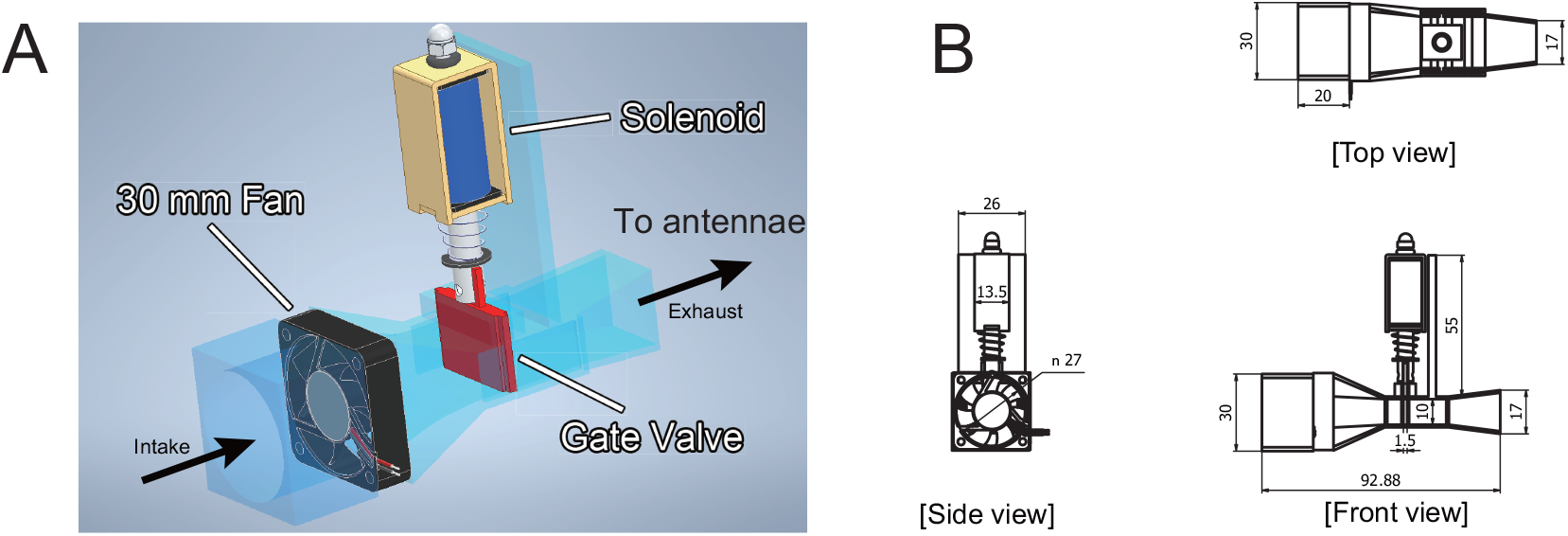
Conceptual diagram and a three-view drawing of the artificial odor acquisition device.

To evaluate the AOA device, we measured the changes in air velocity at the outlet. Most commercially available anemometers have slow time constants and sampling rates of below 1 Hz, which are inadequate for capturing high-frequency variations in air velocity. Therefore, particle image velocimetry (PIV), which measures fluid flow by tracking particle movement using a high-speed camera, was employed to measure the air velocity at the outlet. The setup of the PIV experiment is illustrated in Figure

The distance from the AOA device to the antennae of the silk moth was 25 mm, so we measured the air velocity changes within the red-bordered area, as shown in Figure 3. The laser was aligned with the central axis of the outlet, and fine particles were dispersed from the inlet. The PIV experiment used a high-speed camera (K8, KATOKOKEN Company Ltd., Kanagawa, Japan), laser sheet light source (KLD-V, KATOKOKEN Company Ltd., Kanagawa, Japan), and smoke machine (PS-2005, Dainichi Company Ltd., Niigata, Japan). The fine particles generated by the smoke machine were transferred to a large syringe and drawn in naturally from the inlet side of the AOA device. The camera was configured with a resolution of 800×600 pixels and a frame rate of 800 fps. The air velocity changes were measured at the following frequencies: (1) 1 Hz, (2) 5 Hz, (3) 10 Hz, and (4) 25 Hz.

**Figure 3.**
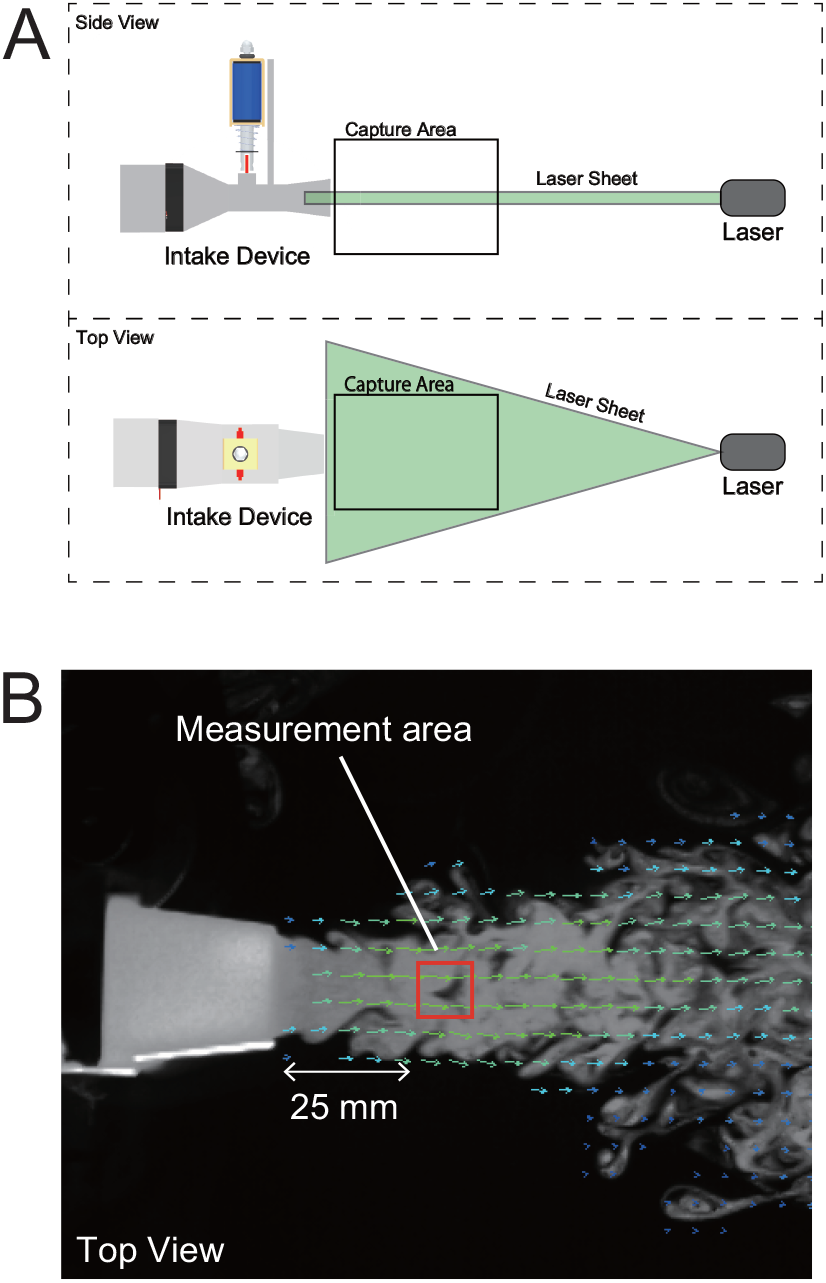
PIV setup for an experiment to evaluate the performance of the AOA device. A: schematic diagram of the experimental setup. B: measurement area for air velocity variation.

The time-series data of the air velocity changes measured in the PIV experiment and the results of the Fast Fourier Transform (FFT) are shown in Figure 4. Examination of the time-series data revealed periodic changes in air velocity for all conditions. We found that there was a clear change in the air velocity up to 10 Hz, but the flow was not completely blocked at 25 Hz. However, FFT analysis of the time-series data revealed distinct peaks at each frequency across all conditions, indicating that odor acquisition can be effectively performed at the desired frequencies. These results demonstrate that the constructed AOA device could achieve an intermittent air intake of up to 25 Hz.

**Figure 4.**
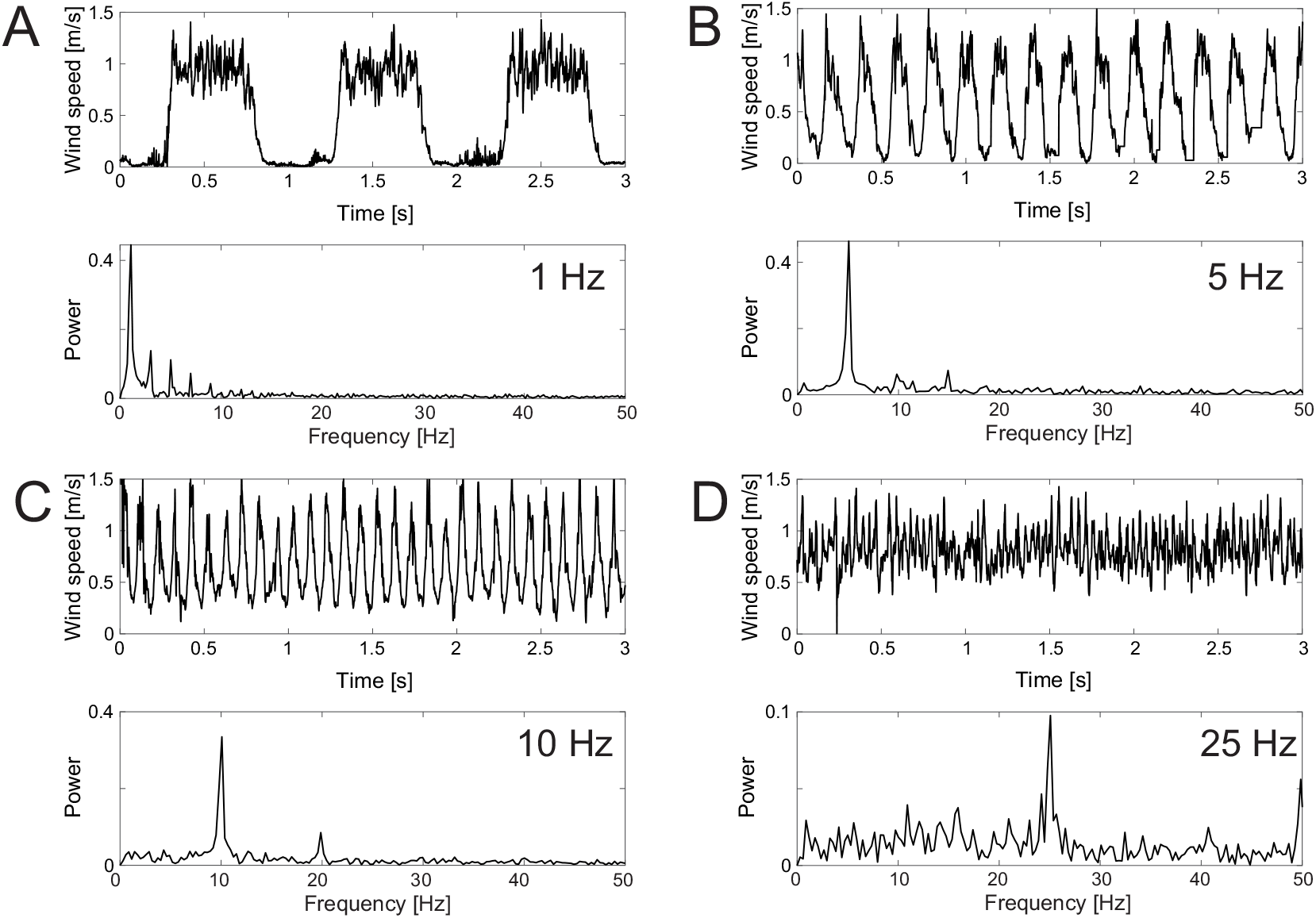
Measurements of air velocity change using PIV, where A-D represent time series data and FFT results at 1, 5, 10, and 25 Hz, respectively.

### 3.3. Insect-mounted robot system

In this study, we performed experiments using an insect-mounted robotic system designed to maneuver a silk moth. The insect-mounted robotic system was equipped with a tethered insect behavior measurement device allowing real-time tracking of an insect ‘s movement speed. The drive system of the robotic system was a differential-wheeled type, enabling it to move in accordance with the measured translational and angular velocities of the silk moth. The robot was controlled based on the movement speed of the moth, and the speeds of the moth and the robot were set to be identical. The behavior measurement device was centered on the robot and used a 60 mm diameter styrofoam sphere. A 40 mm fan was positioned underneath the sphere to levitate it. The sphere guide was fabricated using a 3D printer (Creator3, Flashforge Technology Co., Ltd., Zhejiang, China). Two optical sensors (MA-MA6W, Sanwa Supply Inc., Okayama, Japan) were mounted on the side of the sphere to measure its three degrees of rotational freedom (*θ*_*x*_, *θ*_*y*_, *θ*_*z*_). The optical sensor values were transmitted to the main microcontroller (Arduino Mega) via a microcontroller (Arduino Pro Micro) dedicated to reading the sensor data. The velocity of the silk moth, which was computed using the main microcontroller, was sent to the geared motors. Calibration of the PWM-to-velocity relationship was conducted beforehand because the geared motor was controlled via PWM.

To position the silk moth on top of the styrofoam sphere, a fixture fabricated with a 3D printer was adhered to the back of the moth and fixed to a rod extending from the robot. The fixture was attached with an adhesive (G17, Konishi Co., Ltd., Osaka, Japan) that, according to previous research, does not affect behavior [21].

The entire process of measuring the rotation of the sphere with the optical sensors, converting it to movement speed, and controlling the motor was completed within 10 ms. Previous studies have reported that when the delay between the movement of the moth and the response of the robot exceeds 0.2 seconds, odor plume tracking performance deteriorates [19,20]. Since the processing in this system was completed significantly faster than 0.2 seconds, it was considered that the robotic system did not affect the odor plume tracking performance.

## 4. Odor plume tracking experiment

### 4.1. Experimental design

As illustrated in Figure 5, the odor plume tracking experiments were conducted in a flat open area. At the origin of the experimental field (*x, y*) = (0, 0)[m], bombykol, the sex pheromone of the female silk moth was placed as the odor source. A concentration of 1000 ng of bombykol was applied to a 10 mm diameter filter paper. Air was blown over the filter paper at a flow rate of 4.0 L/min to disperse the odor into the surrounding space. A circulator (YLS-18, Yamazen Corporation, Osaka, Japan) was positioned 0.4 m behind the odor source to facilitate the diffusion of the pheromone. The circulator was adjusted to a wind speed of approximately 0.6 m/s.

**Figure 5.**
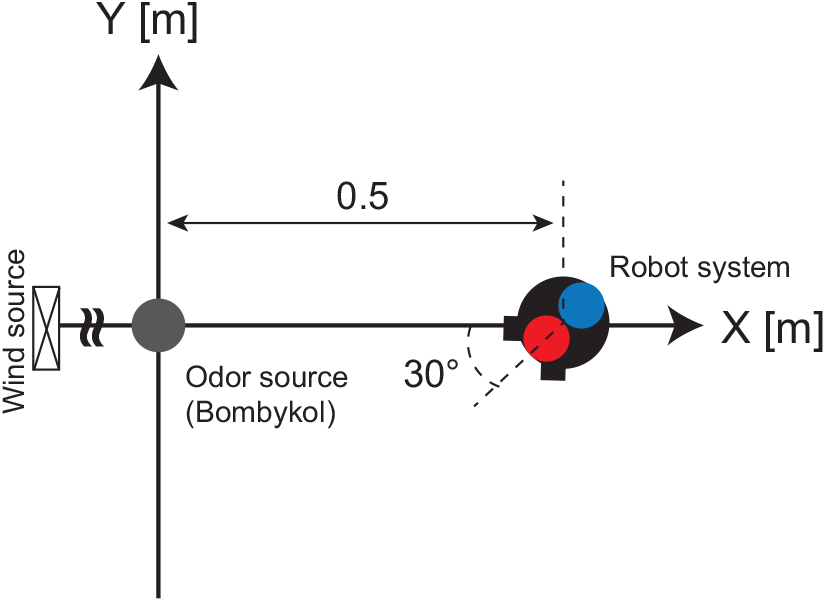
Schematic diagram of the field for odor plume tracking experiment.

The frequency conditions of the AOA device in the odor plume tracking experiment were as follows:

1. Control (gate valve always open)
2. Odor intake frequency at 1Hz
3. Odor intake frequency at 5Hz
4. Odor intake frequency at 10Hz
5. Odor intake frequency at 25Hz

The insect-mounted robotic system was positioned at (*x, y*) = (0.5, 0)[m], and an odor plume tracking experiment was initiated. Localization was defined as successful when the robot reached a position within a radius of 50 mm of the odor source. If the robot failed to achieve localization within 180 s, it was considered a failure. The movements of the robot system were captured using a camera (C910, Logicool Co. Ltd., Lausanne, Switzerland) installed above the field and converted into trajectory data.

The silk moths (Kinshu×Showa; Ehine Sansyu Co. Ltd., Ehime, Japan) used in the experiment were purchased during the pupal stage. The moths had emerged within 2 to 8 days before the experiment. All moths mounted on the robotic system had their wings physically removed to eliminate the possibility of odor intake due to wing flapping. Each AOA device condition was tested using 10 silk moths.

### 4.2 Experimental results

The results of the odor plume tracking experiments are shown in Figures 6. Figure 6A-C represent the localization success rate, localization time, and an index of localization performance derived from the success rate and localization time (SPT, success rate per unit time) [4], calculated using the following equation:

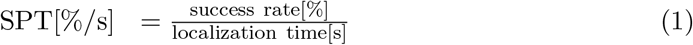

**Figure 6.**
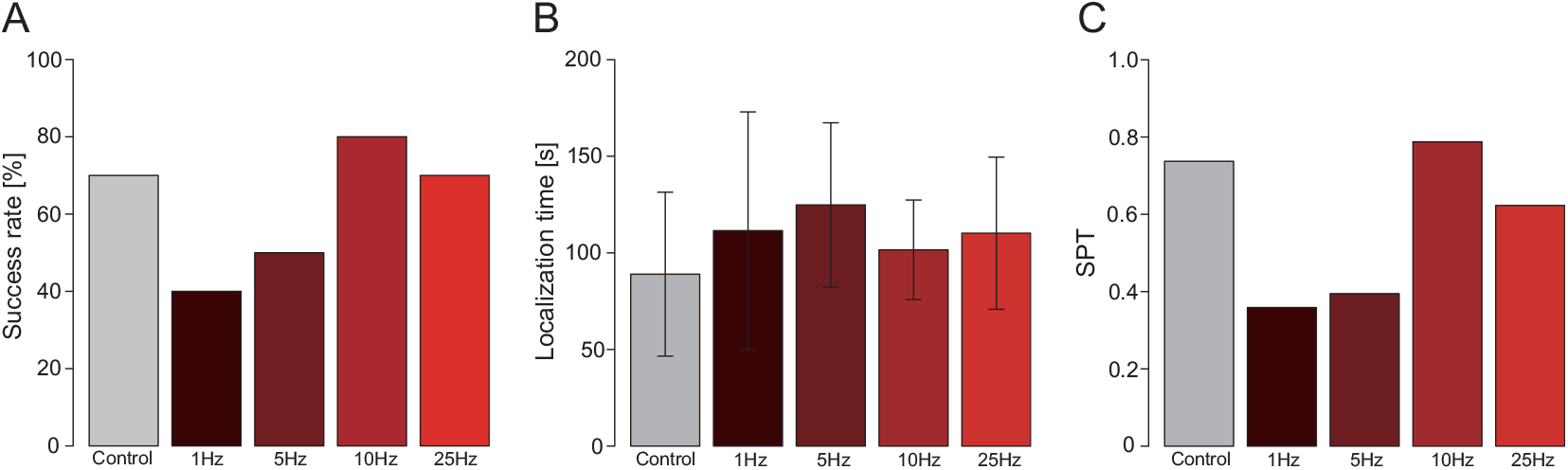
Results of odor plume tracking experiments. A: localization success rate; B: localization time; C: SPT value representing search performance.

Regarding the localization success rate, it is evident that the 10 Hz odor intake condition yields the highest success rate. Conversely, in terms of the localization time, the continuous intake condition (control) achieved the fastest localization. When examining the SPT, which reflects the localization performance, the 10 Hz condition demonstrates the highest value, indicating superior odor plume tracking performance. A comparison of the SPT values of the control and 25 Hz intake conditions revealed almost identical values, with the 10 Hz condition showing slightly better localization performance.

To analyze the behavior of the silk moths under each intake condition, posture angle histograms and localization trajectories are presented in Figure 7. The posture angle histograms in Figure 7A–E indicate the direction in which the moth moved during odor plume tracking, with 0^°^ representing an upwind direction. Blue and black in the trajectory in Figure 7F–J data represent successful and failed localization, respectively. For the 10 Hz condition, where the localization performance is the highest, there is a notable peak near 0^°^, suggesting a more direct movement towards the odor source. In the control condition with continuous odor intake, although there is a high peak in the 0^°^ direction, there are also peaks in other directions, suggesting that continuous strong odor intake does not necessarily result in better performance. Odor intake at a low frequency of 1 Hz caused discontinuities in odor information, which may have elicited movement in different directions. For the 5 Hz condition, although there is an increase in movement toward the odor source, intermittent odor tracking often results in exceeding the time limit. At 25 Hz, the trajectories show behaviors similar to those in the control condition, involving movement in directions other than the upwind direction.

**Figure 7.**
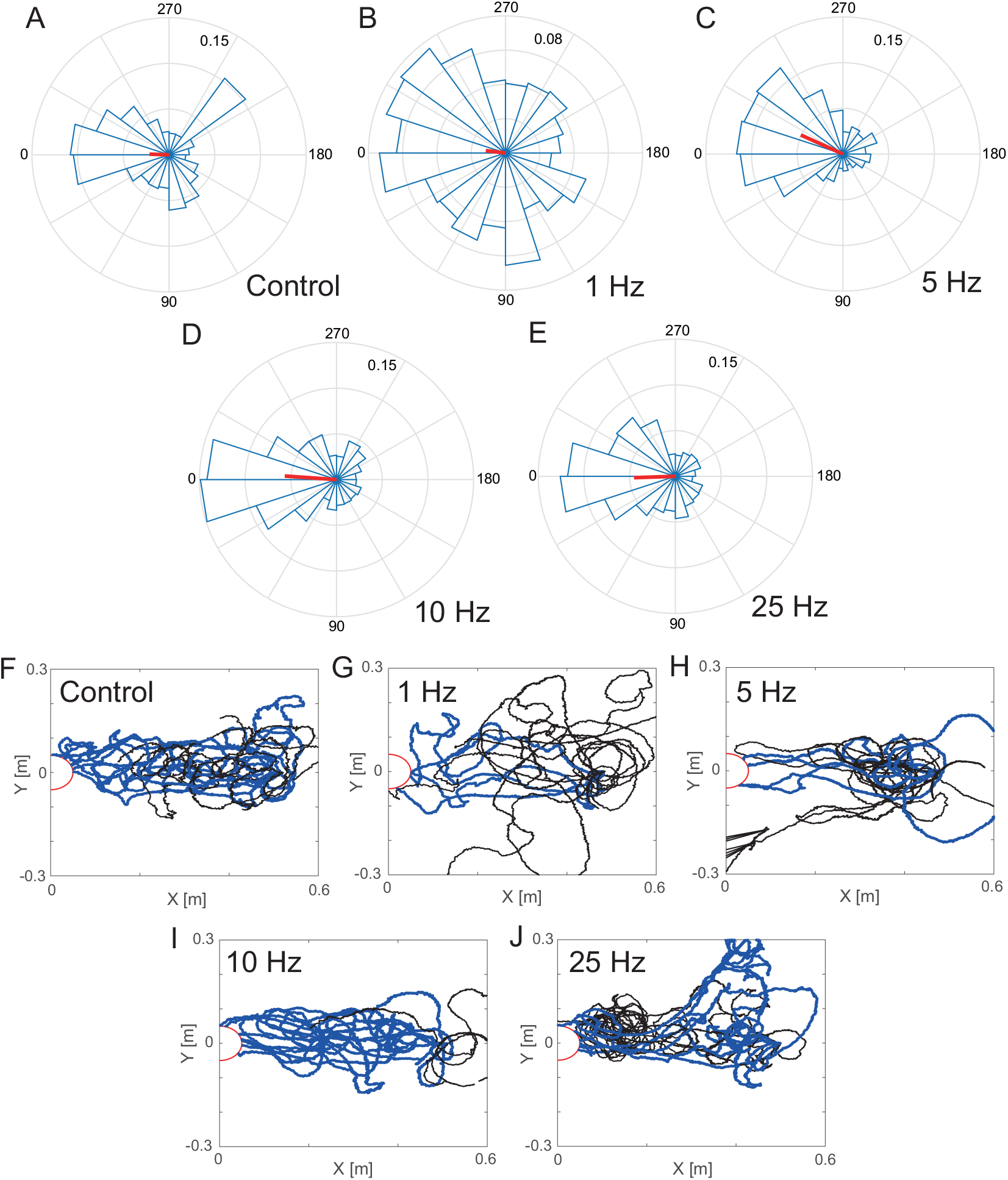
Posture angle change and trajectory during odor plume tracking (*N* =10). A, F: Control; B, G: 1Hz; C, H: 5Hz; D, I: 10Hz; E, J: 25Hz.

At lower frequencies, there was a notable increase in movement in the crosswind direction, resulting in cases in which the silk moth moved out of the odor-existing region. To assess this quantitatively, the extent of movement in the crosswind (Y-axis) direction was represented using a probability density distribution (Figure 8). The area in which the odor emitted from the source was concentrated was investigated in advance using airflow visualization and overlaid on the probability density distribution, as seen in Figure 8. The area of odor presence is depicted in yellow, with the proportion within the yellow region defined as the in-zone ratio and the proportion outside this region defined as the out-zone ratio. A higher in-zone ratio indicates a greater likelihood of continuous odor plume tracking.

**Figure 8.**
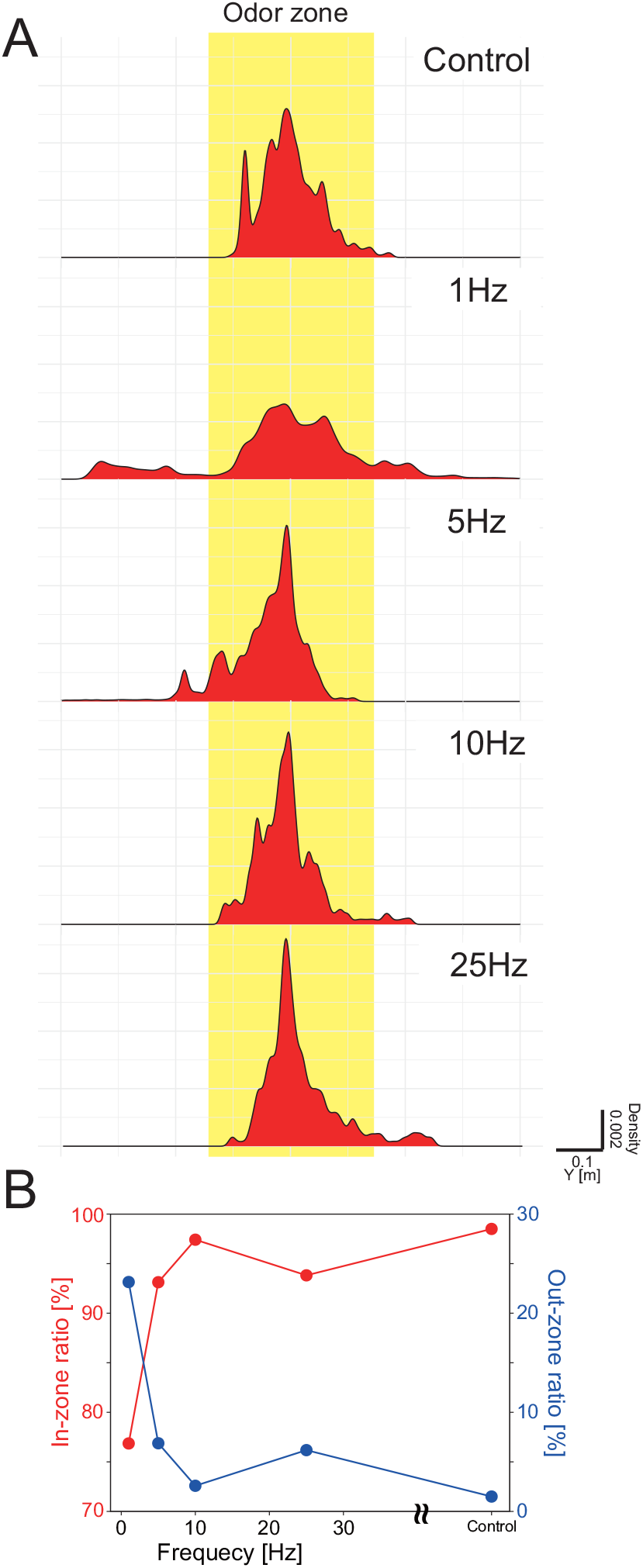
Relationship between trajectory and odor distribution. A: Relationship between the probability density distribution of the trajectory in the Y-axis direction and the odor-existing region for each AOA device condition. B: In-zone ratio and out-zone ratio.

The results under each device AOA condition are shown in Figure 8. It is evident from Figure 8 that the 10 Hz condition achieved the highest in-zone ratio at 98% within each frequency condition. In the cases of 5 and 25 Hz, the in-zone ratio was 93%. Although this was a high in-zone ratio, we found that the odor plume could not always be tracked continuously depending on the situation.

Integrating the AOA device performance results with the odor plume tracking results reveals that intermittent odor intake contributes to improved odor plume tracking performance. Furthermore, modulating the strength of airflow intake and the intermittent frequency affects the behavioral strategy of the moth. Although a constant strong intake enhances the odor plume tracking performance, it sometimes leads to movement in directions different from the odor source, suggesting that intermittent odor intake may suppress such deviations. Therefore, even when implementing a moth-inspired odor plume tracking algorithm, it is important to introduce intermittent odor intake, which is universally performed by animals, or a situation-dependent odor detection method rather than using a sensor system that constantly detects odors.

## 5. Conclusion

In this study, we aimed to investigate the active odor acquisition strategy that enhances the odor plume tracking ability of organisms. In particular, we analyzed the relationship between the intermittent odor entrainment strategy and the localization behavior used by an adult male silk moth. A male silk moth actively engages in odor acquisition through wing flapping; however, intervening in wing flapping to create arbitrary odor acquisition is difficult. Therefore, to investigate these relationships, we introduced engineering tools and employed a robotic system that simulates the wing flapping of silk moths. We designated the device that simulates wing flapping as an artificial odor acquisition (AOA) device and measured changes in the localization behavior of the silk moth in response to varying frequencies of odor acquisition by the AOA device. We found that the difference between air inflow and stoppage in odor attraction was large, and that the odor plume tracking performance was highest at 10 Hz, where odor attraction can be performed at high frequencies. Although continuous strong odor acquisition also improves odor plume tracking performance, it increases the likelihood of movement in directions away from the odor source. This suggests that intermittent odor acquisition at an appropriate frequency helps suppress movement in undesired directions. Thus, to improve odor plume tracking performance, an intermittent odor acquisition strategy at a frequency suited to the localization behavior is crucial. Moreover, it is important to design odor acquisition devices that are suitable for search algorithms when implementing odor plume tracking in robots.

This study focused on odor acquisition during the female localization behavior of male silk moths. Therefore, it cannot be asserted that these results are broadly applicable at this stage, which is a limitation of the present study. Nevertheless, mammals such as dogs and mice exhibit periodic odor-acquisition strategies known as “sniffing” [22], which indicates that intermittent odor-acquisition strategies undoubtedly have some significance.

Future research will involve investigating odor acquisition strategies in other organisms and examining the functional contributions of implementing biological odor acquisition strategies in robotic systems.

## Acknowledgement(s)

The silk moth strains used in the preliminary experiment in this paper were supported by the National BioResource Project (NBRP) of MEXT, Japan.

## Notes on contributor(s)

S.S. and T.M. conceived of and designed the experiments. S.S. and T.M. performed the experiments and analyzed the data, while S.S. and H.O. wrote the paper. N.A. and K.H. contributed to manuscript revision. All authors read and agreed to the published version of the manuscript.

